# Displacement experiments provide evidence for path integration in *Drosophila*

**DOI:** 10.1101/2022.07.22.501185

**Authors:** Anna V. Titova, Benedikt E. Kau, Shir Tibor, Jana Mach, T. Thang Vo-Doan, Matthias Wittlinger, Andrew D. Straw

## Abstract

Like many other animals, insects are capable of returning to previously visited locations using path integration. Recently, *Drosophila* has been added to the list of insects thought capable of this navigational memory. Existing experimental evidence, however, has a potential confound. Here we show that pheromones deposited at the site of reward might enable flies to find previously rewarding locations even without memory. Thus, we designed an experiment to determine if flies can use path integration memory despite potential pheromonal cues by displacing the flies shortly after an optogenetic reward. We found that rewarded flies returned to the location predicted by a memory-based model. Several analyses are consistent with path integration as the mechanism by which flies returned to the reward. We conclude that while pheromones may often be important in fly navigation and must be carefully controlled in future experiments, *Drosophila* may indeed be capable of performing path integration.

## INTRODUCTION

Immediately after a brief taste of sugar, walking blow flies and house flies “dance” – the walking pattern changes into looping trajectories that appear to be a local search, presumably to allow the fly to find additional sugar nearby (Dethier, 1957; White et al., 1984). Recently, such behaviors have been studied in the fruit fly *Drosophila melanogaster*, which also returns to sugar sources using the proposed mechanism of path integration – a memory-based accumulation of distances and directions walked to maintain continually updated information about how to return to a location (Kim and Dickinson, 2017; Brockmann et al., 2018). While many insect species, especially bees, ants and wasps, are known to implement path integration as a form of memory enabling spatial navigation, these recent findings in *Drosophila* are exciting because they suggest the fly neural circuit toolbox may now be used to study the neural basis of navigation in insects. The first steps in this direction have now been made. After optogenetically activating sugar-sensing neurons (Corfas et al., 2019; Behbahani et al., 2021; Lu et al., 2022) and dopaminergic neurons from the PAM cluster (Stern et al., 2019), flies returned to the location of this presumptive reward. The identification of genetically accessible neurons mediating reward in a food search context thus present an entry point into the neural circuitry of foraging in this genetic model organism.

These investigations into *Drosophila* navigation behavior coincide with recent discoveries in the neural basis of navigation in flies, for example, that E-PG neurons in the ellipsoid body neuropil of the central complex integrate angular information to maintain an estimate of self-heading which can be maintained by integration of locomotor turning, visual input, or both (Seelig and Jayaraman, 2015). Recently, Lu et al. (2022) showed that silencing PFNd neurons disrupted performance in an assay designed to test path integration. Thus, it seems that as a field, we are approaching the point where we can address the neural and behavioral basis for path integration in *Drosophila*.

Beyond path integration or other spatial memories, another theoretical possibility that might enable flies to return to sites of previous rewards would be the use of pheromones, chemical cues deposited by oneself or a conspecific. Many ant species deposit pheromone marks and use them to return to important locations. *Drosophila* are known to leave chemical deposits at sites of egg laying (Duménil et al., 2016), and nutritive sugar (Abu et al., 2018) and, at least in some circumstances, flies use such chemicals as a memory-independent cue for spatial navigation (Duménil et al., 2016; Keesey et al., 2016; Lin et al., 2015). Could past behavioral results used to support the hypothesis that flies use path integration have been influenced by a previously unknown contribution from pheromones?

Here we show that flies leave small droplets after optogenetic reward and that naïve flies prefer such locations. Next, inspired by work on path integration in the desert ant *Cataglyphis* (Wehner and Wehner, 1986), we performed displacement experiments to directly test whether *Drosophila* is capable of navigating to a remembered location after being moved away from potential pheromonal cues. Despite a potential ability of flies to use pheromones, the results of our displacement experiments are consistent with flies also being able to use path integration as a behavioral strategy to return to reward.

## METHODS

### Tracking and stimulation software

Real time tracking was based on background-subtraction (Strand Camera, strawlab.org/strand-cam) using a machine vision camera and lens (Ace acA1300-200um, Basler and 12VM412ASIR, Tamron), near infrared illumination (WEPIR1-S1 IR Power LED Star 850nm 1W, WINGER) and filter (Infrared Transmission 87 Filter, Lee). For the pheromone experiment, the stimulation of the emitter flies was performed independent of the flies’ positions. In the displacement experiment during the stimulation period, the LED was turned on if and only if the fly was inside the experimenter-defined and computer-controlled reward zone (RZ). The stages of the experiments were controlled automatically by a Python script that interacted with the tracking software.

### Pheromone experiment

#### Apparatus

For the pheromone experiment, three identical setups were used in parallel. Each experimental setup consisted of a light-tight box with infrared LED illumination, IR sensitive camera for tracking flies in darkness, a red LED for stimulation and a 20 cm diameter walking arena (Figure 1A).

**Figure 1.**
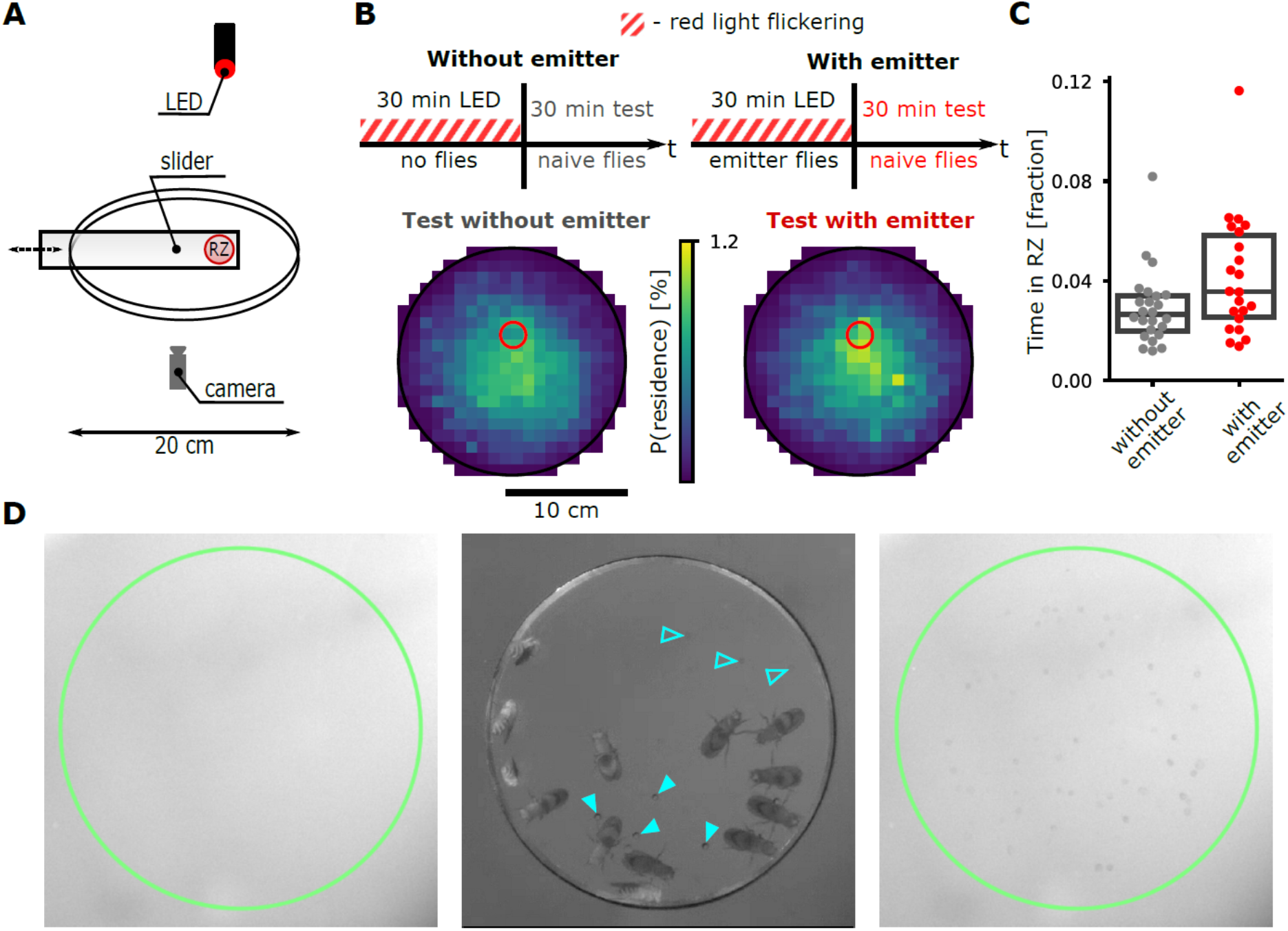
Chemical cues deposited by optogenetically rewarded emitter flies cause accumulation of naïve flies. (A) Experimental setup: a group of emitter flies was put inside a slider with no floor and stood on the arena floor (RZ: reward zone). A group of naïve flies was introduced to the 20 cm diameter arena after the emitter flies were removed via the slider. (B) Top: timeline of the experiment in which emitter flies were illuminated with flickering LED light to activate Gr43a sugar-sensing neurons expressing excitatory optogenetic channels. Bottom: residence probability during the test period for two conditions: without emitter flies (left, n=24 groups), with emitter flies (right, n=22 groups). (C) Fraction of time spent in RZ by naive flies with and without emitter (p = 0.04, 2-tailed Welch’s t-test for unpaired samples) data from B. Every dot represents fraction of time spent in RZ by a group of naive flies (group size: 10-15 individuals). Boxes are from first to third quartile with median indicated as a horizontal line. (D) Magnified view of the reward zone during different stages. Left: arena prior to fly introduction; middle: arena with flies; right: arena after emitter flies removed. The pointers indicate the fresh drops (solid) and the dried out drops (transparent).

The walking arena had circular glass floor and a 3 mm high wall. To prevent the flies from going to the edge of the arena, we built a heat barrier made of aluminum ring thermally glued to a Copper Manganese Nickel alloy heat resistance wire (ISOTAN, 2.5 Ohm/m, 0.5 mm diameter). A coated (Sigmacote, Sigma Aldrich) transparent acrylic lid prevented the flies from leaving or walking on the lid.

The emitter flies were contained in a circular opening in the slider (diameter 20 mm), which was introduced into the arena through the hole in the arena wall. During the slider movement, a flat sheet was put under the slider to prevent the flies contacting arena floor. Between the trials the floor was wiped with ethanol.

The images in Figure 1D were obtained as cropped screenshots from the IR tracking camera. The green circles indicate the object detection area used in the tracking software. To increase the visibility of the released drops, the ImageJ “adjust color balance” tool was applied to the original images. For images without flies, the settings changed were min: 147, max: 245. For the image with flies present, the settings were min: 0, max: 91.

#### Experimental design

The experiment consisted of two stages: emitter stage and test stage (Figure 1B). In the emitter stage, the slider was inside the arena and the stimulation LED was periodically switched (1s on / 2s off). There were two experimental conditions: with emitter --- a group of emitter flies was inside the slider during the first stage; without emitter --- no flies were in the slider. After 30 minutes the slider was removed, naive flies were introduced into the arena (using pipette tips with a cut opening to transfer the flies) and the test stage was recorded for another 30 minutes.

#### Pheromone model of local search in a circular channel

A model of pheromone-mediated behavior was inspired by, and some of the parameters were reproduced from Behbahani et al., 2021. The model consists of a circular linear channel with reward zones and a fly agent that can perform discrete steps (time step 0.5s) in the channel in two directions and release a pheromone drop at any step. The agent is initiated in global search mode, where it continuously walks forward in one direction. When the agent locates the reward (activation zone), it switches to the local search mode. A reward zone can be in one of two states, activated or deactivated. When the agent steps on an activated reward, it performs eating (does not move for 10 time steps), releases a pheromone, chooses a new run length *r*_*rew*_ and continues walking in the same direction for distance *r*_*rew*_ before making a turn. Every time the agent encounters the activated reward, the reward deactivates for a refractory period of 16 time steps. If the agent steps on a released pheromone, it chooses a new run length *r*_*ph*_ which depends on the current odor value of that drop. When the agent finishes a run, it performs a reversal and chooses a new run length *r*_0_ sampled from a wide baseline distribution. The odor value of a pheromone decays over time linearly from 1 to 0 over time *τ*:

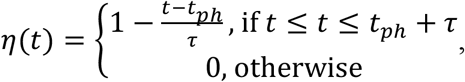

where *t*_*ph*_ is the moment of pheromone drop release. The current run length of an agent is defined by the last action it performed: eating, smelling or reversal, — and does not rely on spatial memory. The run length is chosen from a normal distribution with corresponding parameters.

Reversal:

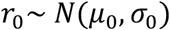

Eating (reward):

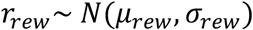

Smelling:

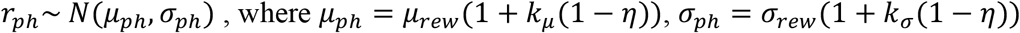

Figure S1 was generated with the following set of parameters: *μ*_0_ = 80, *σ*_0_ = 20, *μ*_*rew*_ = 4.125, *σ*_*rew*_ = 2.625, *k*_*\**_ = 3, *k*_+_ = 3, *τ* = 500. The parameters for eating condition were taken from Behbahani et al., 2021, FR model. Two experiments from the Behbahani study were modeled here: the “circling” experiment and the “multiple rewards” experiment. In the circling experiment model, 6 trials in a row were performed on each fly in a channel with a 26 body length (BL) circumference and the activation zone position is switched between the opposite sides of the channel on every trial. Each trial has a 5 minute activation period (AP) and a 5 minute post-activation (post-AP) period.

The multiple rewards experiment modeling was performed in a bigger 52 BL circumference channel with 3 rewards close to each other (coordinates: 0, ±5 BL), one trial with 5 min AP and 5 min post-AP periods for each simulation.

The run midpoint was defined as the middle between two consecutive reversals, as in Behbahani et al. (2021).

### Displacement experiment

#### Apparatus

The flies walked freely in a circular arena with 60 cm diameter and 20 mm high walls, glass floor and transparent acrylic lid (Figure 2A).

**Figure 2.**
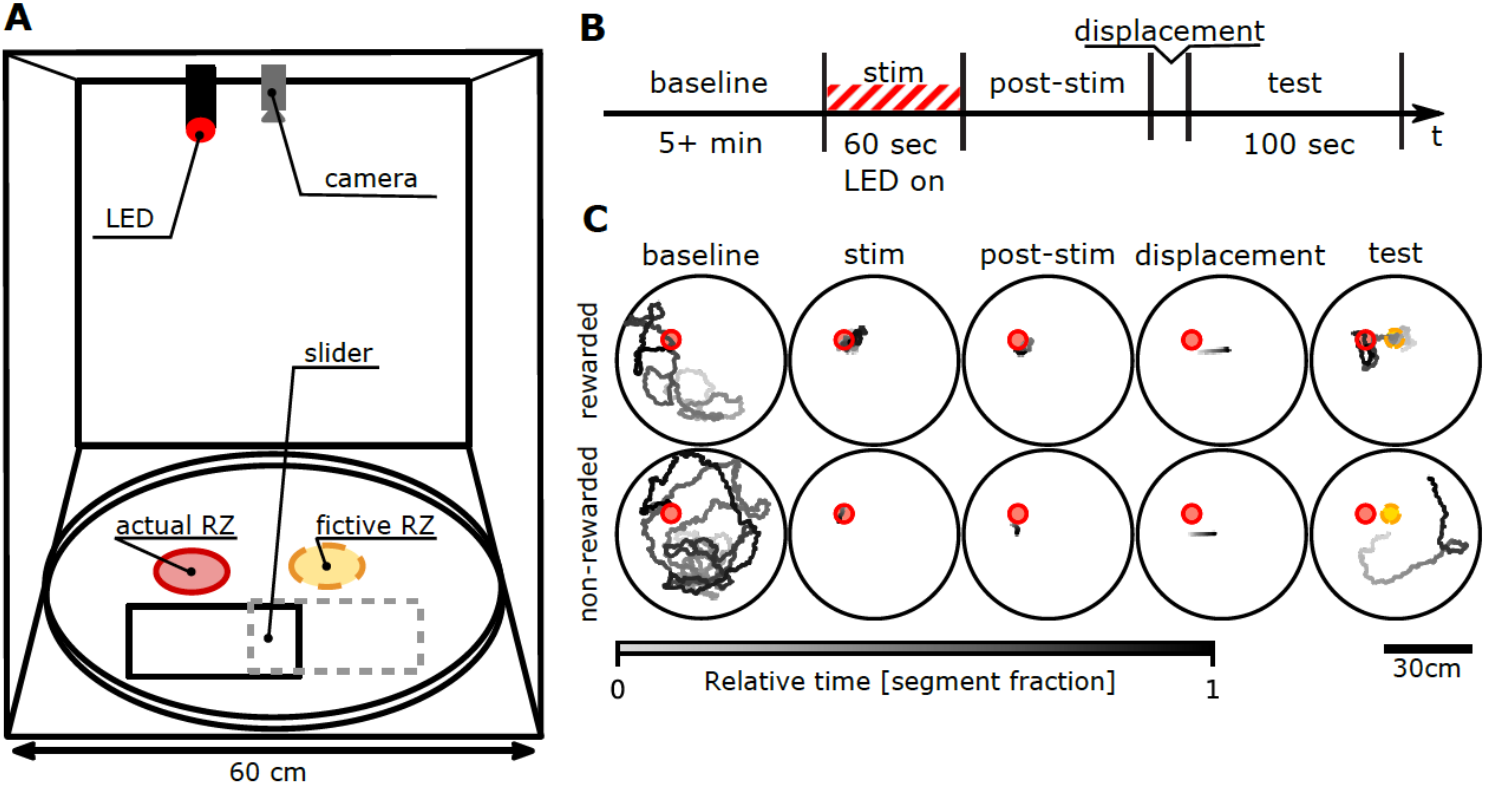
Displacement experiment for freely walking flies tests whether flies use path integration in preference to pheromonal cues. (A) Experimental setup. In 60 cm arena, when the fly left the reward location after the stimulation period and eventually stood on a thin slider, it was manually displaced. (B) Timeline of the displacement experiment for rewarded flies. During the baseline period (“baseline”, 5 – 25 minutes), the fly was introduced to the arena and the reward zone (RZ) was activated after 5 minutes. The stimulation period (“stim”, 60 seconds red light on) was started when the fly found the activated RZ, and the red light was on if and only if the fly was in the RZ. The RZ was deactivated when the fly left the RZ for the first time after the LED on time exceeded 60 seconds. The post-stimulation period (“post-stim”, 3 – 100 seconds), was a wait for the fly to walk onto the slider. In the displacement period (about 3 seconds), the experimenter manually pulled the slider via nylon line with the fly on it. The test period is the first 100 seconds after displacement. For control experiments with non-rewarded flies, the timeline was similar, but the stimulation was not performed. (C) Examples of recorded trajectories during different experimental stages. Top row – rewarded fly, bottom row – non-rewarded fly.

The arena border was made of a bicycle wheel rim, with a Copper Manganese Nickel alloy heat resistance wire (ISOTAN, 10 Ohm/m, 0.25 mm diameter) attached to the outer side, to create a heat barrier preventing flies from going to the edge and climbing the walls. To regulate temperature, small gaps were created by putting washers between the arena floor and the walls, as well as between the walls and the lid. The temperature profile is plotted in Figure S2A.

The entire arena, camera and lens (Ace acA1300-200um, Basler and 12VM412ASIR, Tamron), and LED illumination (generic low power 890 nm LEDs inserted into the spoke holes) were located inside a light-tight cardboard box with black walls and ceiling and white ground (called “experimental chamber” later).

To optogenetically stimulate the experimental flies expressing CsChrimson in targeted neurons, a red LED was mounted above the arena and focused to illuminate a circular part of the arena floor where the flies can be stimulated. The LED stimulation was automatically performed only during the stimulation period when the fly was inside a smaller part of the illuminated area (actual reward zone / RZ), namely a circle with center (x=-9.8 cm, y=6.7 cm) relative to the arena center and 2.8 cm radius. The measured values of the stimulation light intensity are in Figure S2B.

The displacement of the fly was performed using a transparent plastic sheet (A4). It was placed on the arena floor so that the border of the sheet was about 1 cm from the RZ. Two fishing lines were taped to the right side of the sheet, leading outside of the light-tight box where they were pulled by the experimenter in the relocation period.

#### Experimental design

The experimental timeline is illustrated in Figure 2B. The experiment started with a 5-minute baseline period when no optogenetic stimulation was performed and allowed the fly to explore the arena in darkness. Starting from minute 5, the actual reward zone (RZ), would be activated, and as soon as the fly walked inside this circular reward area for the first time, the stimulation period would begin. The red light was turned on when the fly went inside the RZ, and when the fly exited the RZ, the light would turn off. For rewarded flies, the on time was accumulated and when it reached 60 seconds, the stimulation period ended. When the fly exited the RZ, the red light would turn off and would not turn on again. This protocol ensured that all subjects received a minimum cumulative amount of stimulation. Once the fly walked on the plastic sheet slider after stimulation, the slider with the fly on it was displaced by manual pulling on the attached fishing lines. The recording was stopped 10 minutes after the end of stimulation. For analysis, we used the first 100 seconds after displacement.

As a control, a similar experiment was performed without optogenetic reward. Flies of the same genotype were used but the optogenetic LED was never turned on. In this case, the fly had to reach the reward zone and, after that, to go on the slider to be displaced, but no stimulation was performed. Similar to the rewarded condition, for non-rewarded flies, the period of time after a fly first reached the RZ (which was not active) after 5 minutes from the start, to the last time fly left the RZ after spending 0.5 s inside, was called “stimulation” period. The stimulation was controlled automatically by a custom Python script. Note that the displacement slider was pulled manually so the displacement amounts differed across trials (mean length: 92 mm, standard deviation of the mean: 27.8 mm), therefore the fictive RZ location was trial-specific (Figure S3B).

The analysis was performed on all successfully finished trials. We aborted the trial in several conditions: 1) if the fly did not find the RZ in 20 minutes after the RZ activation, 2) if during the stimulation period the fly did not return back to the RZ for more than 2 minutes, 3) if the fly started flying during displacement.

Between the trials, the arena floor and the slider surface were cleaned with 70% ethanol.

### Subject details and environmental conditions

Flies used in the experiments were from a stable stock generated in the Straw lab (generated from BDSC 57636 and BDSC 55136) with genotype *+; Gr43a-Gal4; UAS-CsChrimson::mVenus*.

For the displacement experiment, we used 4–5 day old female flies, starved for 24–30 hours. Before experiments, flies were kept in 25°C, 60% humidity incubator with 12h light/dark cycles (8 a.m. / 8 p.m.). The Gr43a>CsChrimson flies were set on retinal food and kept in the vials wrapped into aluminum foil. The purpose of the foil was to prevent activation of CsChrimson expressing neurons and degradation of retinal in the food due to light exposure. One day before the experiment, the flies were briefly anesthetized at 4°C, sorted by gender and put to starve. During the starvation period, female flies were kept in a foil-wrapped vial with access to water but no food. Approximately 1 hour before start of the experiment, flies were moved into small individual containers (pipette tips) under dim light and put into a dark plastic box. At the experiment start, the fly was taken from the box and placed into the arena. Displacement experiments were conducted in the late afternoon (15:00-20:00). The temperature in the room was kept around 20°C.

For the pheromone experiment, the mixed groups of 4-6 days old male and female flies of the same genotype were used. These flies were raised in the same conditions and starved together in a big vial for at least 24 hours. For the experiment the flies were transferred directly to the arena. The images in Figure 1D were obtained in an independent trial performed on a group of female flies.

### Analysis

Figures were created using FigureFirst package (Lindsay et al., 2017).

#### Data preprocessing

Before the displacement experiment analysis, falsely detected points (outliers) were removed from the raw recordings. The displacement period was identified by manual selection of the corresponding trajectory segment (a relatively straight movement shortly after end of stimulation).

The flies position data from the pheromone experiment were downsampled by time before further analysis.

#### Residency and walking frequency histograms

To illustrate the locational preferences of the flies walking in a circular arena we plotted the residency and walking frequency histograms (Figure 1B; Figure 3A, B).

**Figure 3.**
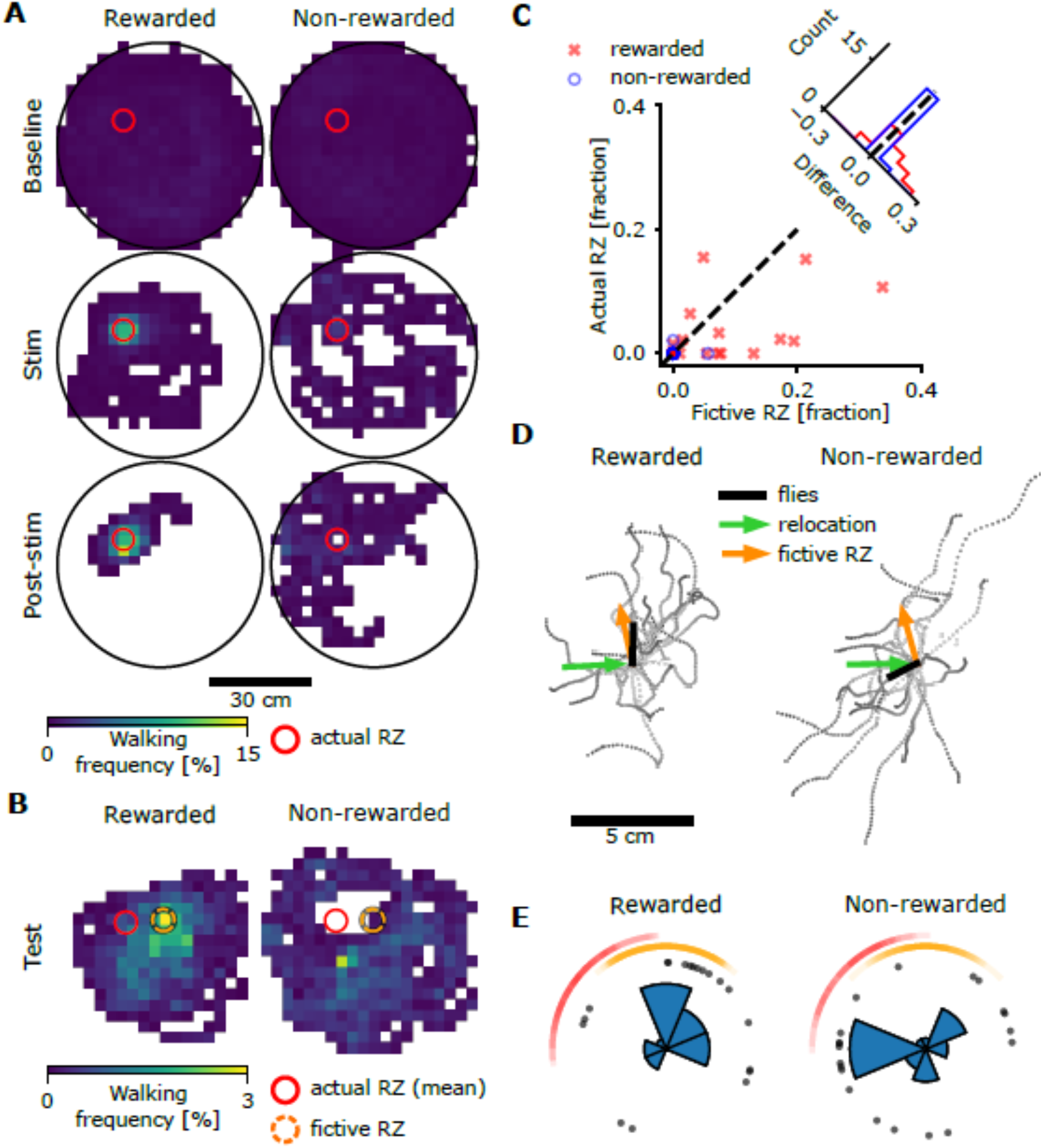
Fly behavior after displacement is consistent with path integration. (A) Walking frequency histograms before displacement for rewarded (n=19) and non-rewarded (n=20) flies: baseline period – light off, stimulation period (Stim) – light on when the fly was inside the reward zone (RZ), post-stimulation period (Post-stim) – light off. (B) Walking frequency histograms during first 100 seconds after displacement (test period) in the fictive reward coordinate system (each individual fly trajectory was aligned to the fictive RZ before making the histogram). (C) Fraction of time spent by the flies inside actual and fictive reward zones during test period (scatter plot) and histograms of corresponding fraction differences. Paired t-test: rewarded p = 0.029, non-rewarded p = 0.572. (D) Trajectories of the flies during first 5 seconds after displacement, shifted to the release point, with indicated trajectory vector average, mean direction of displacement and mean direction to the fictive reward zone. (E) Walking directions in the first 5 seconds after displacement. Each dot represents one fly. Data are rotated such that vertical upwards indicates direction to fictive RZ center. Circular histograms show all flies. The arcs show the range of directions towards the actual (red) and fictive (orange) reward zones.

The residency histogram is a heatmap showing the fractions of trajectory points falling into square-shaped bins of the arena floor (Figure 1B, bin size 1cm x 1cm). To create the residency histograms for the pheromone experiment, we used the data after first five minutes of the test stage. The first five minutes were excluded from the analysis because the flies were much less active during that period of time (data not shown). The resulting heatmap for the experimental condition was created by averaging the heatmaps of individual trials.

For the displacement experiments, we calculated walking frequency histograms (Figure 3A, B) in which points where the flies did not move were removed prior to creating the histogram. The reason is that flies tend to stop in random locations for long periods of time and these stops result in big peaks on the residency histograms, obscuring the local search trajectories during movement (Figure S3A, left column). We did not filter stops for the pheromone experiment because this was a group experiment where the segmentation into individual trajectories would be required but is computationally non-trivial. Due to the larger amount of data, the effect of the individual stops is less corrupting.

To get the ‘walking’ trajectories, a two stage downsampling was performed: first by time, to reduce the noise and number of analyzed points, and then by cumulative distance, with the steps of 0.1 s and 1 mm respectively. This procedure is illustrated in Figure S3A. This two stage downsampling was also used in the distance to RZ analysis (Figure S3D).

#### Fraction of time spent in the area

The fraction of time spent by the fly in actual or fictive reward zone (Figure 3E) was calculated based on the trajectories downsampled by time with step 0.1 s.

#### Distance to reward zone

The distance to reward zone was defined as the distance from the fly to the center of the reward zone. The distributions of distances to actual and fictive reward zones (Figure S3D) were plotted based on the 2-stage downsampled trajectories (described in Methods: Walking frequency histograms).

#### Translation of axes

The fictive reward location was unique for every individual fly, because the displacement direction and length were not the same. To analyze the statistics of fly locomotion relative to the fictive reward zone, the post-displacement trajectories were shifted in a way that the fictive reward location had similar coordinates for all flies: a new coordinate system with origin at fictive reward location was introduced. A similar approach was used to address the question of how the action of displacement influences fly walking direction: the axes were translated to the release point (end of displacement) and the walking directions from this point were compared in different conditions.

#### Statistical comparisons

To analyze the walking directions after displacement (Figure 3E) we performed a Rayleigh test for unimodal deviations of circular uniformity independently for rewarded and non-rewarded conditions, using the test.rayleigh function from Python package pycircstat (https://github.com/circstat/pycircstat/). The mean direction was calculated as angular component of vector average of all directions represented as unit vectors. For other null hypothesis significance testing, we used scipy.stats functions.

#### Intersection score

As a measurement of intersection between fly trajectories just before (entry trajectory) and just after (exit trajectory) displacement (Figure S3F, G) we used the length of the longest intersecting segment of these trajectories. For every point of the exit trajectory, we calculated the distance to the entry trajectory as the minimum of all distances to the points of the entry trajectory. The intersection score was defined as the length of the longest segment of the exit trajectory, all points in which were closer than the intersection threshold t (2.5 mm) to the entry trajectory.

## RESULTS

### Visitation to particular spatial locations can be mediated by pheromones

To address the potential role of pheromones in a fly navigation task, we checked if the previous presence of optogenetically rewarded flies, called emitter flies, could affect the spatial behavior of naïve flies introduced to the arena later (Figure 1). Therefore, we compared the behavior of naïve flies in two conditions: a condition with emitter flies and a control condition without emitter flies which was otherwise treated identically. All flies (naïve and emitter) expressed the light gated cation channel CsChrimson using *Gr43a-GAL4* to target sugar sensing neurons and thus would experience the rewarding taste of sugar when exposed to red light. During the initial 30 minute period, with or without a group of emitter flies, a flickering red LED illuminated the reward zone (RZ). After this initial reward period, a slider was removed from the arena which, in the condition with emitter flies, also removed them from the arena (Figure 1A). Naïve flies were introduced and the arena was kept in darkness for 30 minutes while their position was tracked using infrared illumination and camera. Histograms of naïve fly position show greater residence close to the reward zone for the trials in which emitter flies had previously been optogenetically rewarded (Figure 1B). The fractions of time each test group of flies spent inside the reward zone (Figure 1C) were statistically significantly different between the two conditions (p=0.04, unpaired two-tailed Welch’s t-test, N emitter=22 groups, N no_emitter=24 groups). Photographs before, during and after emitter flies were rewarded (Figure 1D) show droplets deposited by the flies. These results suggest that pheromones are sufficient to cause naïve flies to accumulate at the site of reward and could be used as a navigational strategy available to *Drosophila* when performing optogenetic reward experiments.

These results do not allow us to distinguish whether accumulation of naïve flies depends on the emitter flies having been rewarded, as we tested only rewarded flies. Regardless, these results highlight the possibility that chemical cues deposited by emitter flies, even in a potentially reward-independent way, could cause accumulation of naïve flies. Previous studies which used fly accumulation as evidence for path integration generally have not controlled for the potential effect of fly-emitted pheromones (Corfas et al., 2019; Kim and Dickinson, 2017; Stern et al., 2019), and therefore we argue that they cannot be used as strong evidence that flies indeed are capable of idiothetic path integration.

One line of evidence used to argue flies perform path integration comes from recent experiments in which flies were given an optogenetic reward, again by activation of sugar-sensing neurons, in a specific location along a circular channel (Behbahani et al., 2021). In that work, repeated visits to the rewarded location, especially after multiple cycles of walking around the circular channel, were interpreted as evidence of path integration. To address whether these results might be theoretically explained by a pheromone-based mechanism rather than a path-integration mechanism, we created an agent-based model performing the same task (Figure S1). In our model, a simulated fly agent can emit pheromones which decay in intensity over time but this agent has no ability to remember the locations of previous rewards. The statistics of the agent behavior in this model were similar to those observed in real flies and we tested simulated behavior in two experimental designs based on those of Behbahani et al. In the first design, reward location alternated between two opposite sides of the circular channel, and we observed a distribution of ‘run midpoints’ (see Methods for definition) similar to that measured experimentally (Figure S1A-C). The second experimental design used the same model with three reward locations, and we got a distribution of run midpoints similar to those both of the path integration model of Behbahani et al. and their experimental results (Figure S1D-F). In these simulations, we designed the model structure and manually adjusted parameters to check if a pheromone-based agent model could approximate the behavioral data and found that indeed it could. While a closer match to behavioral data could be made by fitting experimental data with optimization algorithms, this is not necessary to establish the fundamental point that a pheromone-based mechanism would be sufficient to reproduce several lines of evidence used to argue that flies are capable of path integration. Our experiments (Figure 1), those of others (Duménil et al., 2016; Lin et al., 2015) and our modeling (Figure S1) show that pheromones need to be considered as a potential mechanism by which flies return to rewarding locations in the environment. Furthermore, experimental data that was argued to support the path integration hypothesis (Behbahani et al., 2021; Kim and Dickinson, 2017) have not adequately excluded the possibility that the results may include a contribution from a pheromone-based mechanism.

### Displacement experiments provide evidence for path integration in *Drosophila*

Now we ask whether flies can use path integration, independent of possible use of pheromones, to return to rewarding locations. To do so, we implemented a procedure in which flies walking in a circular arena in darkness were automatically rewarded when they entered a 5.6 cm diameter circular reward zone (RZ) via optogenetic stimulation with a computer-controlled LED. Following the reward period, we waited for them to walk onto a thin slider and then moved them from the RZ vicinity – and thus away from any emitted chemicals or other sensory cues physically linked to this location – and displaced them by several centimeters (Figure 2, details in Methods). If flies use path integration as a dominant mechanism to return to the reward location, they should return to the ‘fictive RZ’, the remembered location of the reward which would now be decoupled from the ‘actual RZ’ at which they physically received the reward. Alternatively, if flies used pheromones or other cues physically linked to the site of actual reward, they should not particularly visit the fictive RZ but rather the actual RZ. By displacing the flies from the actual RZ, we reduced the potential effect of pheromones and, if flies can use multiple mechanisms, sought to increase the relative importance of path integration mediated returns.

As expected, rewarded flies did spend a substantial duration of time walking in the RZ during optogenetic stimulation and immediately afterwards, prior to displacement. Control flies, which did not receive the optogenetic reward, did not spend substantial time there (Figure 3A). The key question is whether, after the displacement, the rewarded but not control flies would walk disproportionately towards, and in vicinity of, the fictive RZ. Histograms of walking frequency suggest that rewarded flies indeed spent substantially more time in the fictive RZ compared to the actual RZ, consistent with the path integration hypothesis (Figure 3B, Figure S3C). Notably, the peak of walking activity in the fictive RZ in the rewarded flies suggests that flies remember both the distance and direction to the rewarding location. In the 100 second test period, 13 of 19 rewarded flies visited the fictive RZ, compared to one of 20 control flies (two proportion z-test: z = 4.13, p = 3.7e-5). Thus, optogenetic reward led to a higher probability of visiting the fictive RZ after displacement. (The actual RZ was also visited more frequently by the rewarded flies than by the non-rewarded flies – 9/19 rewarded vs. 1/20 non-rewarded, z = 3.03, p = 0.002.) Quantification shows that rewarded flies spent statistically significantly more time in the fictive RZ than the actual RZ, whereas control flies did not (paired t-test: rewarded p = 0.029, control p = 0.572, Figure 3C).

If flies are using path integration, one may predict that their initial motion after displacement would be in the direction of the remembered reward, for example by following an accumulated vector to reward location. We thus examined the initial direction of movement after displacement. Rewarded flies move on average in the direction of the fictive RZ after displacement (Rayleigh test for uniformity: p = 0.018, z = 3.894; mean angular deviation from the fictive RZ center: 28.9°), whereas control flies did not move in any consistent direction (Rayleigh test: p = 0.352, z = 1.057; mean direction relative to the fictive RZ center: −92.0°) (Figure 3D,E, Figure S3E).

Theoretically, the results so far are consistent with a model in which rewarded flies emit a pheromone trail along their path and, after displacement, return along this chemical trail. We ranked the flies by the amount of intersection between their trajectories just before and just after displacement (2 cm radius from the displacement points; Figure S3F,G). Only four trajectories had an intersection value of 1 cm or greater and appear as if such route following may have occurred (Figure S3F). Two such trajectories were from rewarded flies and two from non-rewarded flies. Additional quantification did not reveal differences between the rewarded and control groups (Figure S3G). These results are inconsistent with the pheromone trail hypothesis, as are the results of a previous study which addressed a similar question (Brockmann et al., 2018).

## DISCUSSION

Using experiments and modeling, we showed that flies may emit pheromones that could be used to guide the fly to return to a previously rewarded location. Such chemical cues may be a simple yet effective strategy to return to rewarding locations and our data suggest this possibility must be carefully controlled in studies of path integration behavior. Taking this potential confound into account, we performed a displacement experiment which tested the possibility that flies use path integration to return to a remembered location of previously experienced optogenetic reward. Our results support the hypothesis that flies are able to use path integration to maintain an estimate of distance and direction to the location of a prior reward. These results thus add to the growing evidence that *Drosophila* can perform path integration but suggest careful delineation of pheromone-mediated from memory-mediated mechanisms will be required as spatial navigation behaviors are further investigated.

Future studies could investigate the chemical composition of the deposits seen here and how they relate to deposits known from existing work (Abu et al., 2018; Duménil et al., 2016; Keesey et al., 2016; Lin et al., 2015). By storing information about the environment in pheromones, the ultimate effect – of returning to a previously rewarding location – may be achieved with a substantially simpler computational strategy.

It will be interesting to study how path integration interacts with other navigational strategies and how additional sensory input, such as visual cues, which were excluded by our experimental design here, interact with the path integration abilities we studied here. To pursue the neural basis of these behaviors, the neuro-genetic tools available to *Drosophila* researchers could be used to investigate the neural circuits involved in path integration. Several recent studies have taken advantage of *Drosophila* fixed rigidly to record neural activity in the central complex – a key neural substrate for navigation behaviors – during walking or flying behavior (Fisher et al., 2019; Giraldo et al., 2018; Green and Maimon, 2018; Green et al., 2017; Kim et al., 2017; Lu et al., 2022; Seelig and Jayaraman, 2015; Turner-Evans et al., 2017). Although restraining the animal facilitates brain imaging and electrophysiology, it may also limit behavioral performance as it changes biomechanics in addition to disrupting the multi-model input an animal would normally receive as it walks or flies freely. Preventing natural, free movement is known to disrupt normal visuo-motor coordination in flight (Stowers et al., 2017) and we predict that tethering also may alter navigational abilities. The evidence in support of path integration we presented here and that in the literature comes from experiments in freely walking flies (Behbahani et al., 2021; Brockmann et al., 2018; Corfas et al., 2019; Kim and Dickinson, 2017; Stern et al., 2019). One recent such study argued that intact PFNd neurons are required for path integration in a free walking assay (Lu et al., 2022). In addition to path integration, the navigational capabilities of freely walking *Drosophila* include returning to rewarding locations using vision (Foucaud et al., 2010; Ofstad et al., 2011) and using a working memory for visual landmarks (Neuser et al., 2008). An open question is thus whether path integration or other natural navigation behaviors can be replicated in a tethered behavioral apparatus. More generally, efforts to bridge the gap between the ability to perform physiological recording from tethered flies and the ability to study behaviors of freely moving flies will be useful as we seek to investigate the circuits for navigation in *Drosophila*. Emerging and future work will seek to increase the realism of tethered experiments, for example by increasing the sophistication of the feedback using virtual reality in tethered animals (Haberkern et al., 2019). Complementary efforts to perform calcium imaging of neuronal activity in freely walking flies attempt to bridge the gap from the other direction (Grover et al., 2020). A combination of neuronal silencing and activation in freely behaving animals in addition to physiological experiments in tethered animals also looks promising (Lu et al., 2022).

*Drosophila* may be an ideal system to investigate the relevant neural circuits and, ultimately, the biophysical implementation of path integration in one species of insect. Many species of insects are capable of path integration, with the list recently expanded to include *Scarabaeus galenus* dung beetle (Dacke et al., 2020). Path integration is a well-established component of the suite of navigational strategies used by Hymenoptera (Stone et al., 2017; Wehner and Wehner, 1986; Wittlinger et al., 2006) and has now been described in a non-insect member of the Arthropoda, *Neogonodactylus oerstedii* mantis shrimp (Patel and Cronin, 2020). Thus, the question arises if capability of performing path integration may have evolved in ancient ancestors of the Arthropoda and if the behavioral abilities and basic neural substrate may have been conserved since then. If so, *Drosophila* will serve as a useful model system across the insects and beyond.

## ACKNOWLEDGEMENTS

We thank the mechanical and electrical workshops of the Institute of Biology I, University of Freiburg. This work was funded by SmartSmart 2 Fellowship from the Bernstein Network Computational Neuroscience to AVT and DFG grant STR 1357/6-1 to ADS. We thank Bloomington Drosophila Stock Center for providing the transgenic fly lines used here.

## SUPPLEMENTAL FIGURES

**Figure S1.**
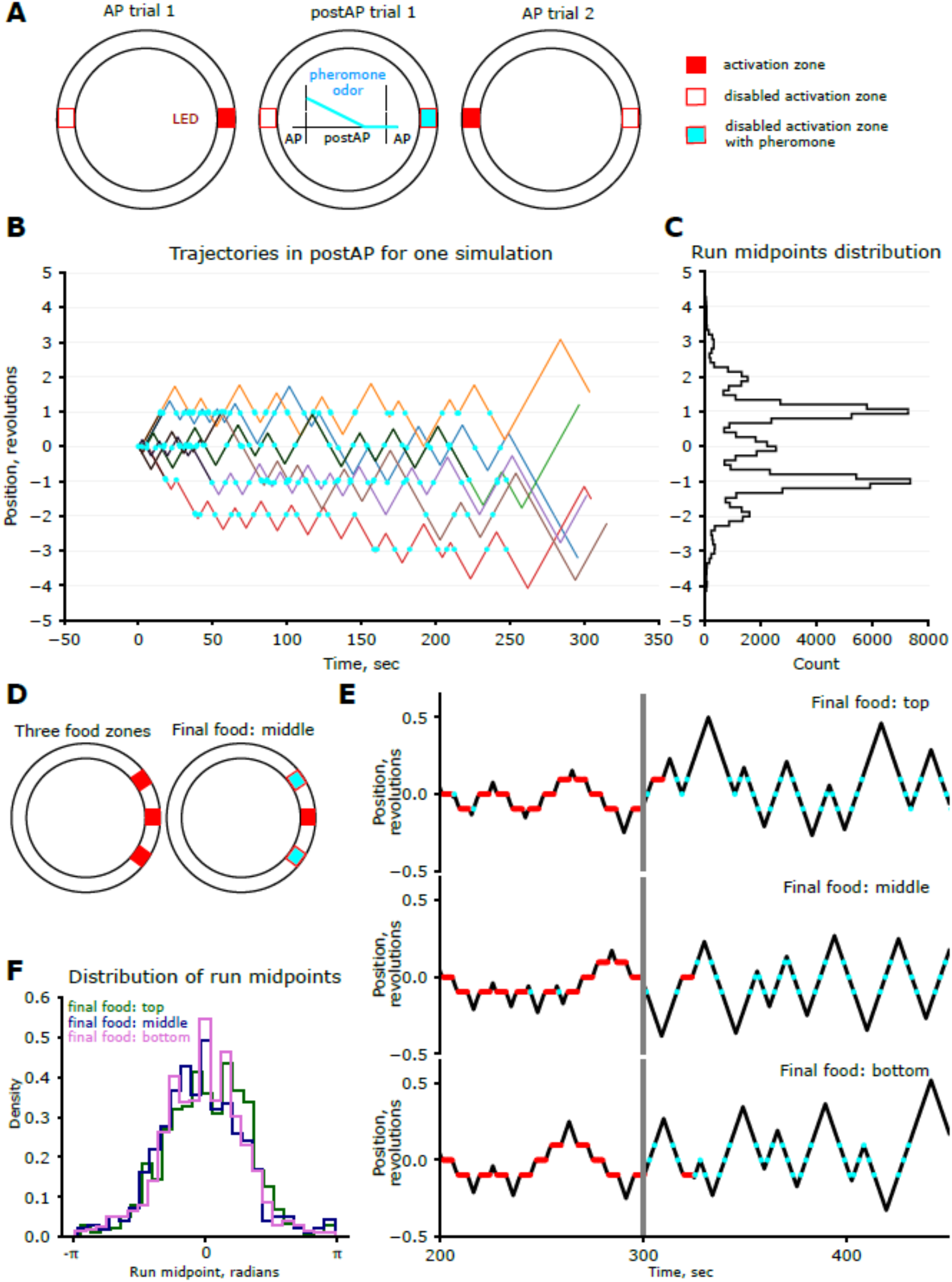
The pheromone model of Drosophila navigation in a circular channel. (A) Schematic of the small annular arena (26 body lengths circumference) with alternating activation zones. After being rewarded in an activation zone, the agent releases a pheromone with a decaying odor value (compare with Behbahani et al. 2021, Fig. 2). (B) Example pre-return (black) and post-return (colorful) trajectories of pheromone model simulation in a small arena. Time 0 corresponds to the last eating and is highlighted in red. The position is relative to the last reward. The moments where the agent feels the pheromone odor are highlighted in cyan. (C) The histogram of run midpoints of the post-return trajectories from 1000 simulations. (D) Schematic of the bigger annular arena (52 BL) with three food zones (compare with Behbahani et al. 2021, Fig. 6). For the last activation one of three food zones is used. The deactivated zones will have a pheromone signal. (E) Example trajectories of the pheromone model simulations for different final food locations. (F) The distribution of run midpoints after activation for different final food locations. Only trials where the agent visited all three food zones in last two AP runs were used. Total number of simulations: 1000 for each condition.

**Figure S2.**
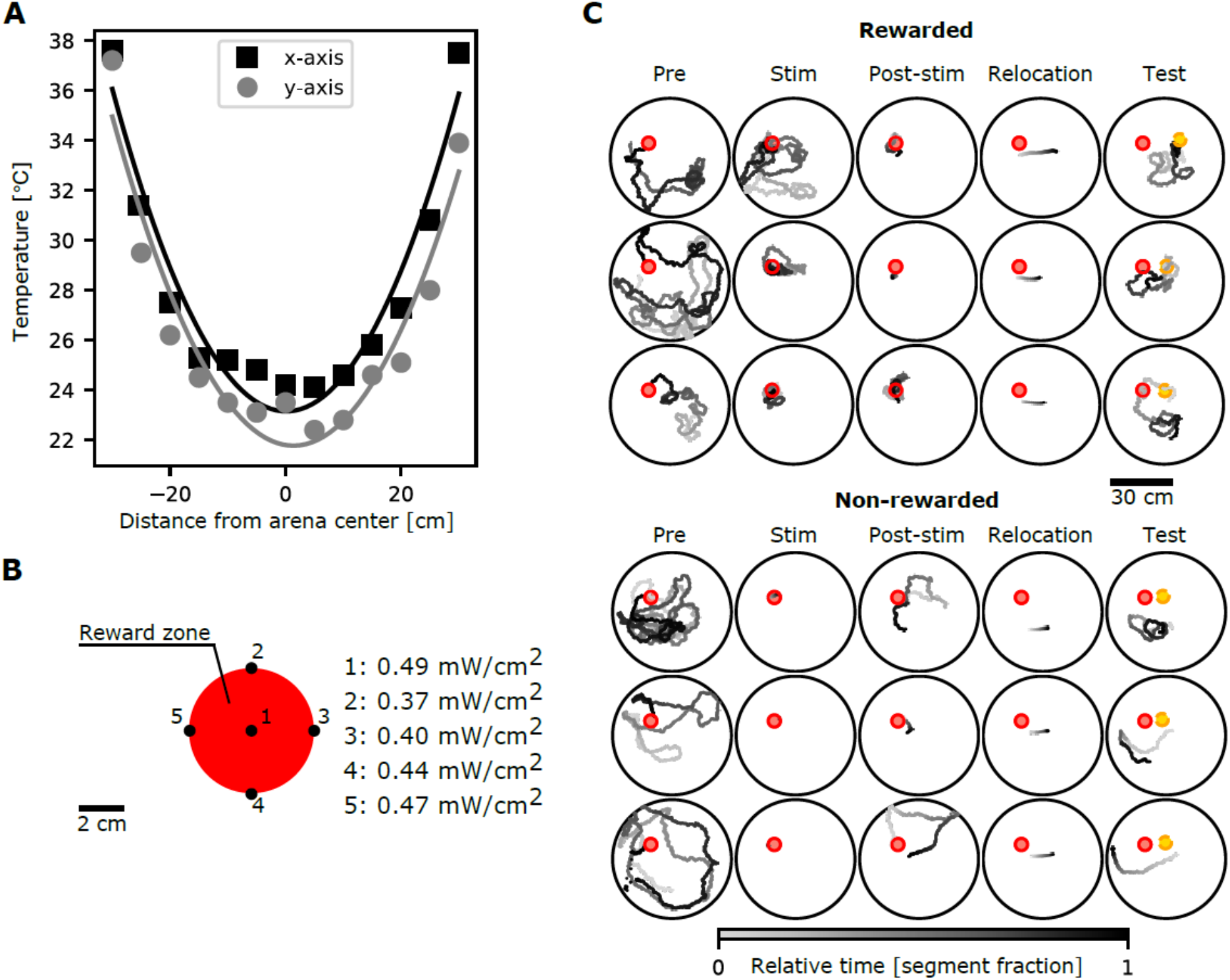
Characterization of optogenetic stimulus. (A) Temperature profile along two axes with 2-degree polynomial approximations. (B) Values of stimulated light intensity in some points in the reward zone. (C) Trajectories of three rewarded and three non-rewarded flies during different experiment stages. Each row is from a single fly.

**Figure S3.**
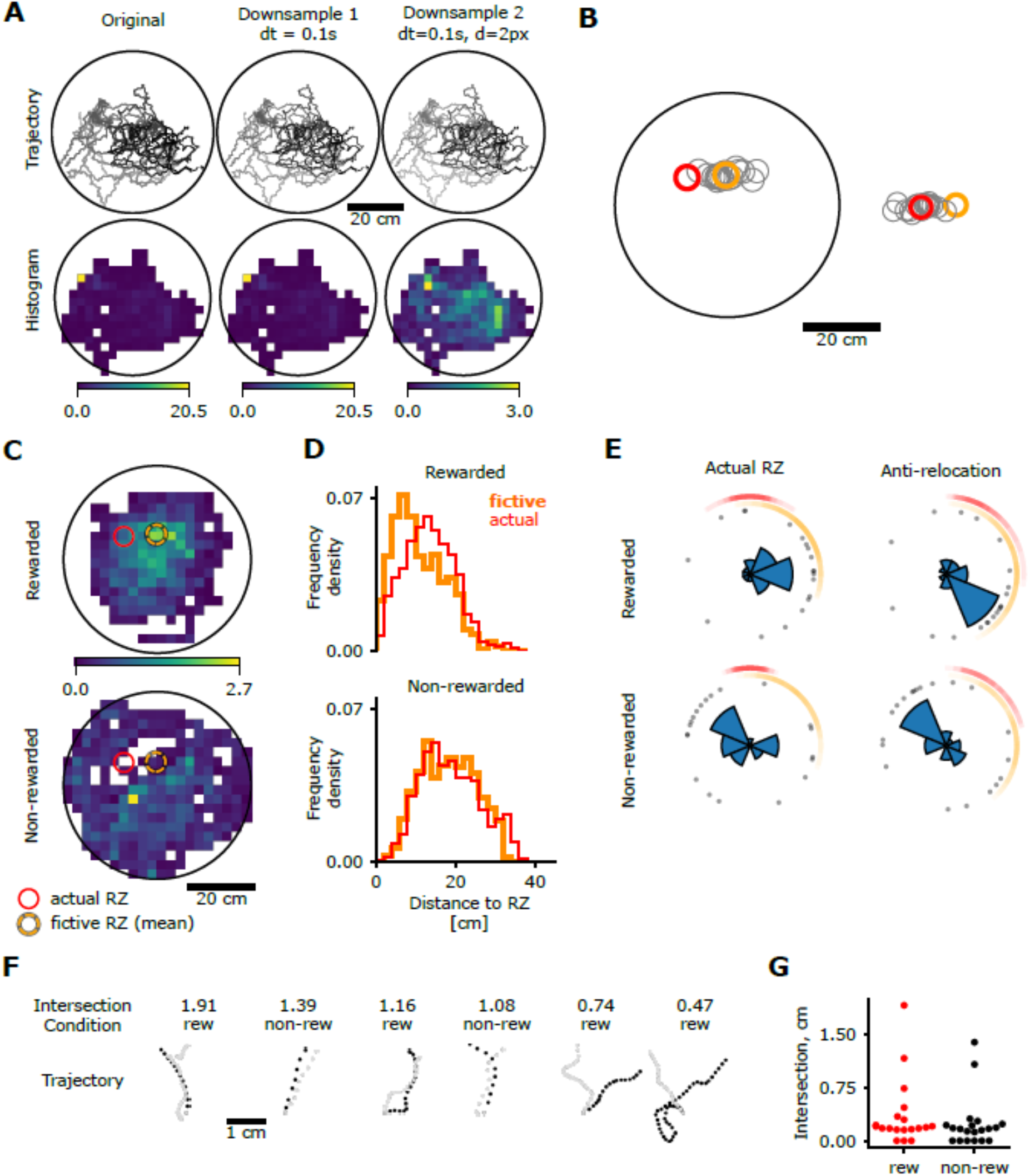
Displacement experiment analysis. (A) 2-stage downsampling to calculate walking locations from residency locations. Left to right: original --- frame rate 100 fps; stage 1 --- by time (step 0.1 s), stage 2 --- by distance (step 1 mm). (B) Coordinate transform: left --- original coordinate system, right --- fictive RZ coordinate system. Reward zones for all trajectories are indicated in gray, mean reward zones in color (actual: red, fictive: orange). (C) Walking frequency histograms during test period in the original coordinate system. Same data aligned to fictive RZ: Figure 3B. (D) Distributions of distances from all points visited by the flies during test period to fictive (orange) and actual (red) RZ. (E) Walking directions during first 5 seconds after displacement. Each dot represents one fly. The arcs show the range of directions towards the actual (red) and fictive (orange) reward zones. Compared to plots in Figure 3D, these are aligned to the actual RZ (left) and displacement direction (right). (F) Trajectories immediately before (in gray) and after displacement (in black) with highest intersection scores. Rewarded flies: “rew”, non-rewarded flies: “non-rew”. (G) Amount of intersection between trajectories immediately before and after displacement, comparison between rewarded (n=19) and non-rewarded (n=20) flies. Each dot represents one fly. Mann-Whitney U rank test: p = 0.230.

